# Effects of inhibitors on Hsp90’s conformational dynamics, cochaperone and client interactions

**DOI:** 10.1101/255190

**Authors:** Sonja Schmid, Markus Götz, Thorsten Hugel

## Abstract

The molecular chaperone and heat-shock protein Hsp90 has become a central target in anti-cancer therapy. Nevertheless, the effect of Hsp90 inhibition is still not understood at the molecular level, preventing a truly rational drug design. Here we report on the effect of the most prominent drug candidates, namely radicicol, geldanamycin, derivatives of purine and novobiocin, on Hsp90’s characteristic conformational dynamics and the binding of three interaction partners. Unexpectedly, the global opening and closing transitions are hardly affected by Hsp90 inhibitors. Instead, the conformational equilibrium, as well as the associated kinetic rate constants remain almost untouched. Moreover, we find no significant changes in the binding of the cochaperones Aha1 and p23 nor of the model substrate Δ131Δ. This holds true for both, competitive and allosteric inhibitors. Therefore, direct inhibition mechanisms, affecting only one molecular interaction, are unlikely. Based on our results, we speculate that the inhibitory action observed *in vivo* is caused by a combination of subtle effects, which can be used in the search for novel Hsp90 inhibition mechanisms.

## Introduction

The molecular chaperone and heat-shock protein Hsp90 is a metabolic hub (***Taipale et al., 2010***). It is involved in all *Six Hallmarks of Cancer* (***Blagg and Kerr, 2006***). Due to this exceptional role, Hsp90 has become a central target in a broad range of anti-cancer therapies. Numerous studies have been undertaken, both by academia and industry, to develop Hsp90 inhibition strategies (***Biamonte et al., 2010; Neckers and Workman, 2012; Sidera and Patsavoudi, 2014; Taldone et al., 2014; Zuehlke et al., 2018***). Nevertheless, the molecular basis of the observed therapeutic effects is still unknown, although it is essential for rational drug design. Mainly four classes of Hsp90 inhibitors have been investigated as anti-cancer drug candidates so far. Three of them bind to Hsp90’s unusual, N-terminal ATP binding site, a rare Bergerat fold (***Dutta and Inouye, 2000***). These are derivatives of geldanamycin (***DeBoer et al., 1970***), radicicol (***Delmotte and Delmotte-Plaquee, 1953; Khandelwal et al., 2016***) and purine (e.g. PU-H71 (***Immormino et al., 2006***)). They are competitive inhibitors, which suggests ATPase inhibition as the mechanism, but this assumption has not been demonstrated yet. In addition, there is an allosteric inhibitor class: the novobiocin derivatives, sometimes called novologues (***Garg et al., 2016; Marcu et al., 2000a; Matts et al., 2011***) (e.g. KU-32 (***Anyika et al., 2016***)), which targets Hsp90’s C-terminal domain. While the class of purine derivatives originates from *in silico* studies (***Chiosis et al., 2001; Immormino et al., 2006***), all the other main classes are natural product derivatives found by screening (***Khandelwal et al., 2016***). Although *in vivo* studies report measureable effects of these inhibitors, there is currently little *molecular* understanding of the inhibition mechanism of these drug candidates.

It is generally accepted that any mechanistic hypothesis must stand both, *in vivo* and *in vitro* testing. In an attempt to provide a better *molecular* understanding of anti-cancer drug candidates targeting Hsp90, and to complement the existing *in vivo* results, we report how well-known small molecular inhibitors affect other important molecular observables in Hsp90’s functional cycle *in vitro*.

First we probe the characteristic conformational changes between a v-shaped open conformation with dissociated N-domains, and a compact closed conformation where the three domains (N-terminal, middle, C-terminal) of the homodimer form inter-monomer contacts with their equivalent counterparts (***Ali et al., 2006; Hellenkamp et al., 2017; Southworth and Agard, 2008***). It is commonly assumed that these characteristic conformational changes are rate-limiting for Hsp90’s function, involving ATP hydrolysis, co-chaperone interaction and finally client processing (***Prodromou, 2012; Sattin et al., 2015; Schopf et al., 2017***).

First, we use single molecule Förster resonance energy transfer (smFRET), which is perfectly suited to reveal conformational dynamics that are usually hidden by ensemble averaging (***Ha, 2001; Mickler et al., 2009***). We attach one donor and one acceptor dye to specific residues of either Hsp90 monomer (Figure 1). The sensitivity of FRET on inter-dye distance changes enables us to distinguish open and closed conformations of Hsp90 and it allows us to record conformational changes in real-time with a total internal reflection fluorescence microscope (TIRFM). We provide a quantitative description of these kinetics, which became available by our recently developed single molecule analysis of complex kinetic sequences (SMACKS) (***Schmid et al., 2016***). Altogether, this single molecule approach is very sensitive to drug induced changes in the relative population of open and closed conformations, and also to changes in transition kinetics between those.

Second, we investigate the *in vitro* effect of the inhibitors on the binding of two well characterized cochaperones, namely p23 (***McLaughlin et al., 2006***) and Aha1 (***Li et al., 2013***) using fluorescence anisotropy.

Third, we examine a possible interference of the inhibitors with substrate binding: the established model client Δ131Δ (***Street et al., 2011***) binds between the M-domains of the Hsp90 dimer (***Schopf et al., 2017***), which is representative for many other clients.

Altogether we find that there is not a single straightforward inhibition mechanism for any of the major Hsp90 inhibitor classes, indicating that they might rather act on diverse features of the highly dynamic chaperone system.

**Figure 1.**
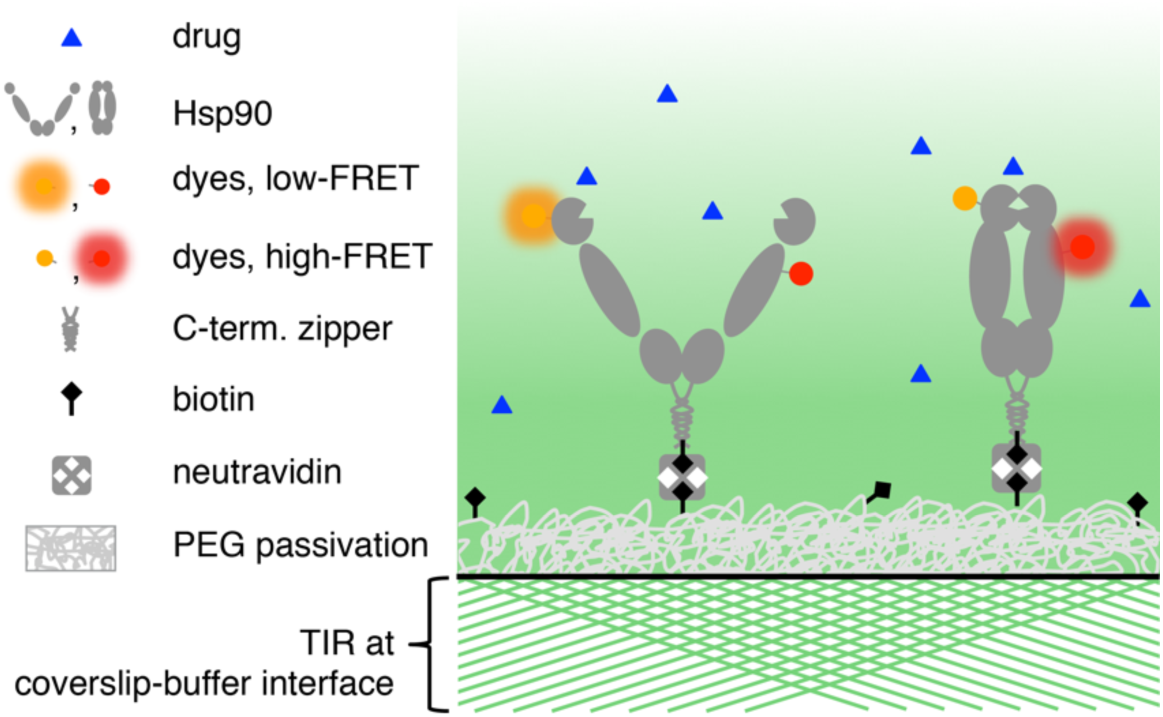
Illustration of the smFRET experiment revealing the effect of drug candidates on Hsp90’s conformational dynamics. Single molecule FRET between the attached fluorescent dyes allows us to distinguish open and closed conformations of Hsp90. To follow one molecule for minutes, the dimers are stabilized by a C-terminal zipper motif and immobilized on a polyethylene glycol (PEG) passivated coverslip using biotin-neutravidin coupling. Fluorescence intensities of individual dyes are recorded by total internal reflection (TIR) fluorescence microscopy. Schematic laser rays are depicted as green lines. The evanescent excitation intensity is shown in fading green. See example data in Figure 2A.

## Results

This study includes one lead compound of each main class of Hsp90 inhibitors, namely geldanamycin, radicicol, the purine derivative PU-H71 (***Immormino et al., 2006; Rodina et al., 2007***) and novobiocin inspired KU-32 (***Anyika et al., 2016***) at saturating concentrations (see Materials and Methods section). Figure 2A shows example fluorescence traces obtained from individual fluorescently labeled Hsp90 dimers in the presence of one of these inhibitors, respectively. Specific conformational transitions are observed as an anti-correlated change in donor and acceptor fluorescence. In the closed conformation of Hsp90, both dyes are close to each other (approx. 53 Å) leading to high acceptor fluorescence and low donor fluorescence, due to efficient FRET. The opposite is the case in the open conformation, where the dyes are further apart (approx. 92 Å), causing low acceptor fluorescence and high donor fluorescence. The information from a total of over 600 molecules is combined in the FRET efficiency histograms in Figure 2B. We have previously shown that yeast Hsp90’s prevalent conformation - under many conditions including saturating ADP as well as ATP - is an open, v-shaped one (***Hellenkamp et al., 2017***), leading to low FRET efficiencies in the described smFRET experiment. In contrast, in the presence of the non-hydrolysable ATP analogue AMP-PNP, Hsp90 occurs mainly in the globally closed conformation (cf. crystal structure 2cg9 (***Ali et al., 2006***)).

**Figure 2.**
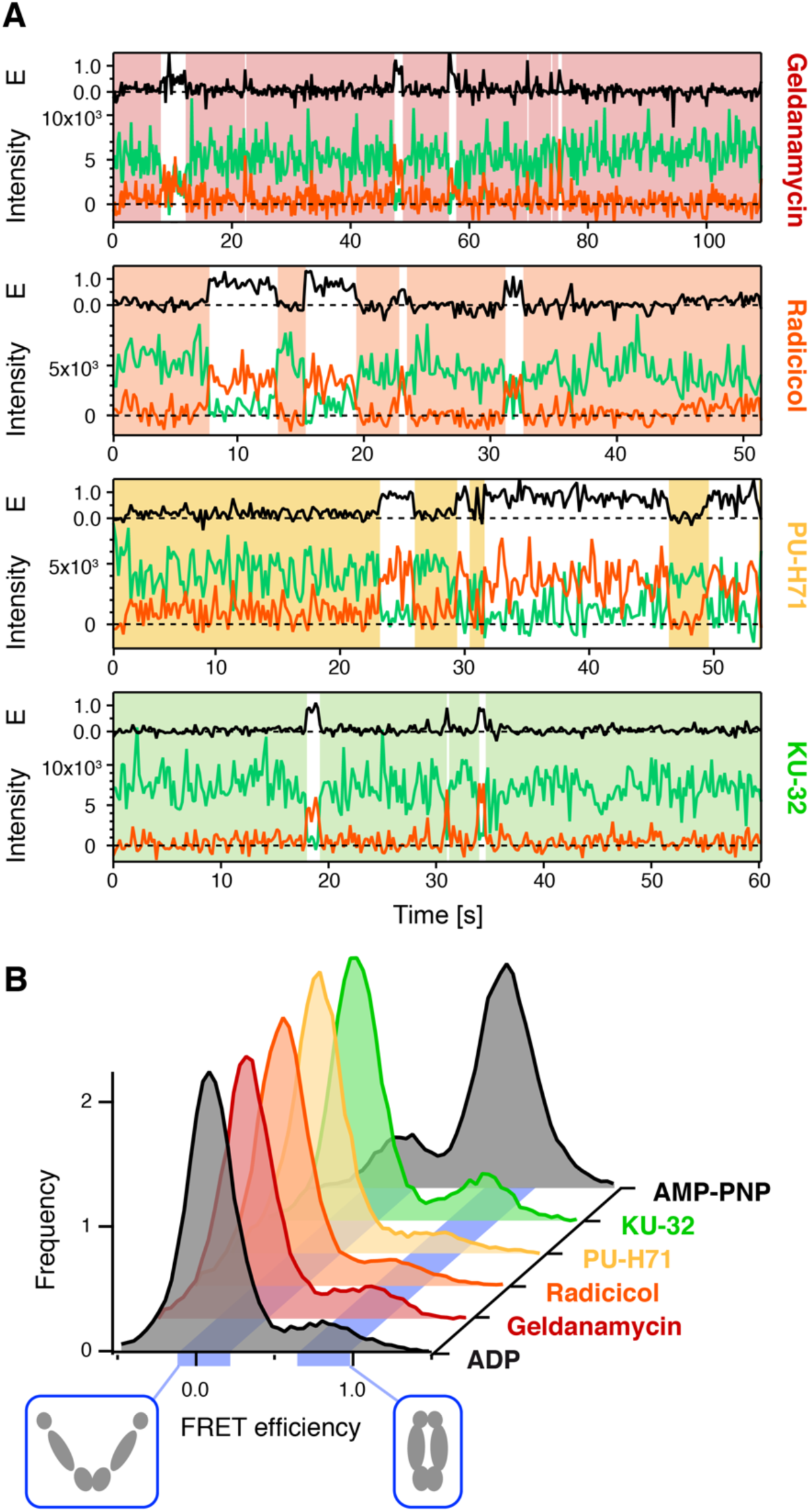
(**A**) smFRET trajectories show conformational dynamics in the presence of the indicated inhibitor. Fluorescence intensities of individual FRET donor and acceptor dyes coupled to Hsp90 are shown as green and orange lines, respectively. The resulting FRET efficiency (E) is shown in black. Hsp90’s closed and open conformations are indicated as white and colored overlays, respectively, given by the Viterbi path (**Schmid et al., 2016**). (**B**) FRET histograms (normalized to unity) show that open conformations (low FRET) prevail under all conditions, except for the non-hydrolysable nucleotide analogue AMP-PNP, which stabilizes the closed conformation. Blue ribbons highlight the expected FRET efficiencies of the indicated open and closed conformations. n(ADP)=107, n(geldanamycin)=108, n(radicicol)=142, n(PU-H71)=123, n(KU-32)=65, n(AMP-PNP)=104.

Interestingly, Figure 2B shows that clearly none of the four lead compounds provoked a shift similar to AMP-PNP. While individual example traces (Figure 2A) show some statistical variation, neither the competitive inhibitors nor the allosteric KU-32 changed the equilibrium distribution considerably. Therefore, a systematic shift in the conformational equilibrium directly caused by the inhibitors is most unlikely to cause the inhibitory effects observed *in vivo*. Obviously, identical equilibrium distributions can be caused by multiple sets of kinetic rate constants, i.e. the conformational *kinetics* could still vary. Therefore, we analyzed the observed kinetics more thoroughly using the software SMACKS (***Schmid et al., 2016***).

As shown in Figure 3A, a 4-state model with 3 links was found for Hsp90 in the presence of each of the four inhibitors, no matter from which class they were. The states 0/1 represent open conformations; states 2/3 denote closed conformations. Globally open or globally closed conformations may differ in local structural arrangements. Although not all four conformations are directly distinguishable in terms of FRET efficiency, the observed kinetic behavior implies a 4-state model (***Schmid et al., 2016***). We generally observe the fastest transition rates between the short-lived states 1 and 2, whereas states 0 and 3 represent longer-lived but less frequently accessed states. For either inhibitor, state 0 is most populated followed by state 3, 1 and 2. The corresponding quantitative rate constants are displayed in Figure 3B. The uncertainty of the rate constants is reported as their 95% confidence interval. In the presence of AMP-PNP all transition rates differ significantly from the ADP case. The decreased opening and increased closing rate constants perfectly explain the drastic shift in the FRET histogram upon the addition of AMP-PNP in Figure 2B. In contrast, the effect induced by the inhibitors is small: while some significant differences are observed, these lie only marginally outside the confidence interval. There is also no systematic difference between the N-and C-terminal inhibitors. Altogether, the observed small effects of the inhibitors on Hsp90’s global conformational changes, cannot on their own explain the inhibitory effects found *in vivo*.

**Figure 3.**
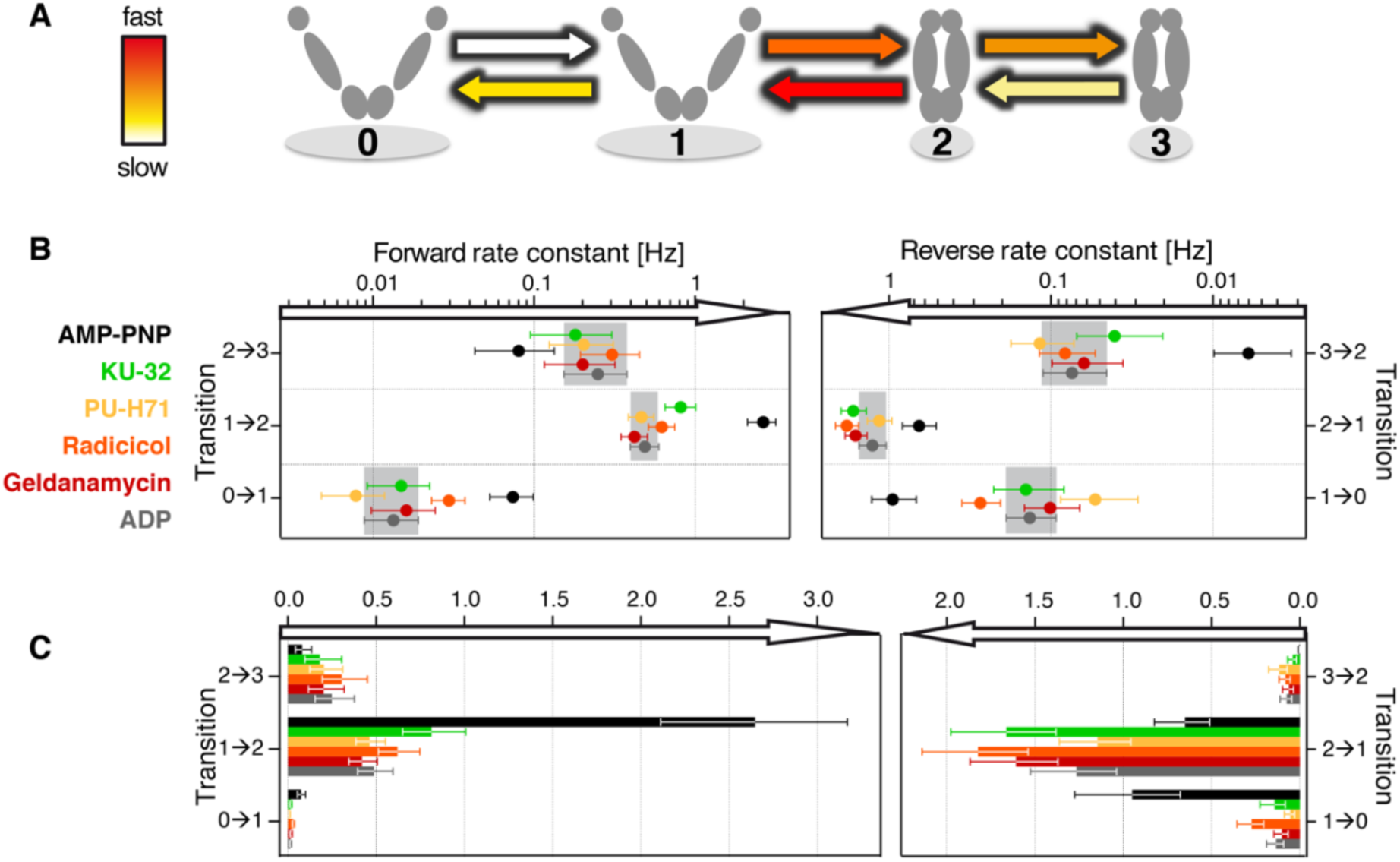
Kinetic model of Hsp90’s conformational dynamics. (**A**) Schematic consensus model under all tested inhibitor conditions, as well as ADP. (**B**) Quantitative rate constants and confidence intervals of the transitions in (**A**). Gray areas represent the 95% confidence intervals in the presence of ADP. Significantly different rate constants were found with AMP-PNP (black), but none of the inhibitors changed the kinetics in a similar way. (**C**) Same data as in (B) plotted in a linear scale.

Next we investigate the effect of the inhibitors on Hsp90’s interaction with cochaperones and a model client. Figure 4 shows the fluorescence anisotropy of the binding partners in absence and presence of the inhibitors. 10 μM Hsp90 and saturating amounts of the inhibitors were added to 200 nM Aha1 (fluorescently labeled at position 85), 400 nM p23 (labeled at position 2) or 400 nM Δ131Δ (labeled at position 16). As a control 2% (vol:vol) DMSO was added. In every single case the DMSO control shows larger or similar effects compared to the inhibitor. Therefore, it is unlikely that impaired binding of Hsp90 to these cochaperones or the client, is the reason for the inhibitory effects found *in vivo*. Again, this holds for N-and C-terminal inhibitors. Note that the binding sites for these three investigated binding partners are at complementary positions. Aha1 and p23 mainly bind the closed state of Hsp90 at the N and M domains, while Δ131Δ binds mainly the open state in between the M domains.

**Figure 4.**
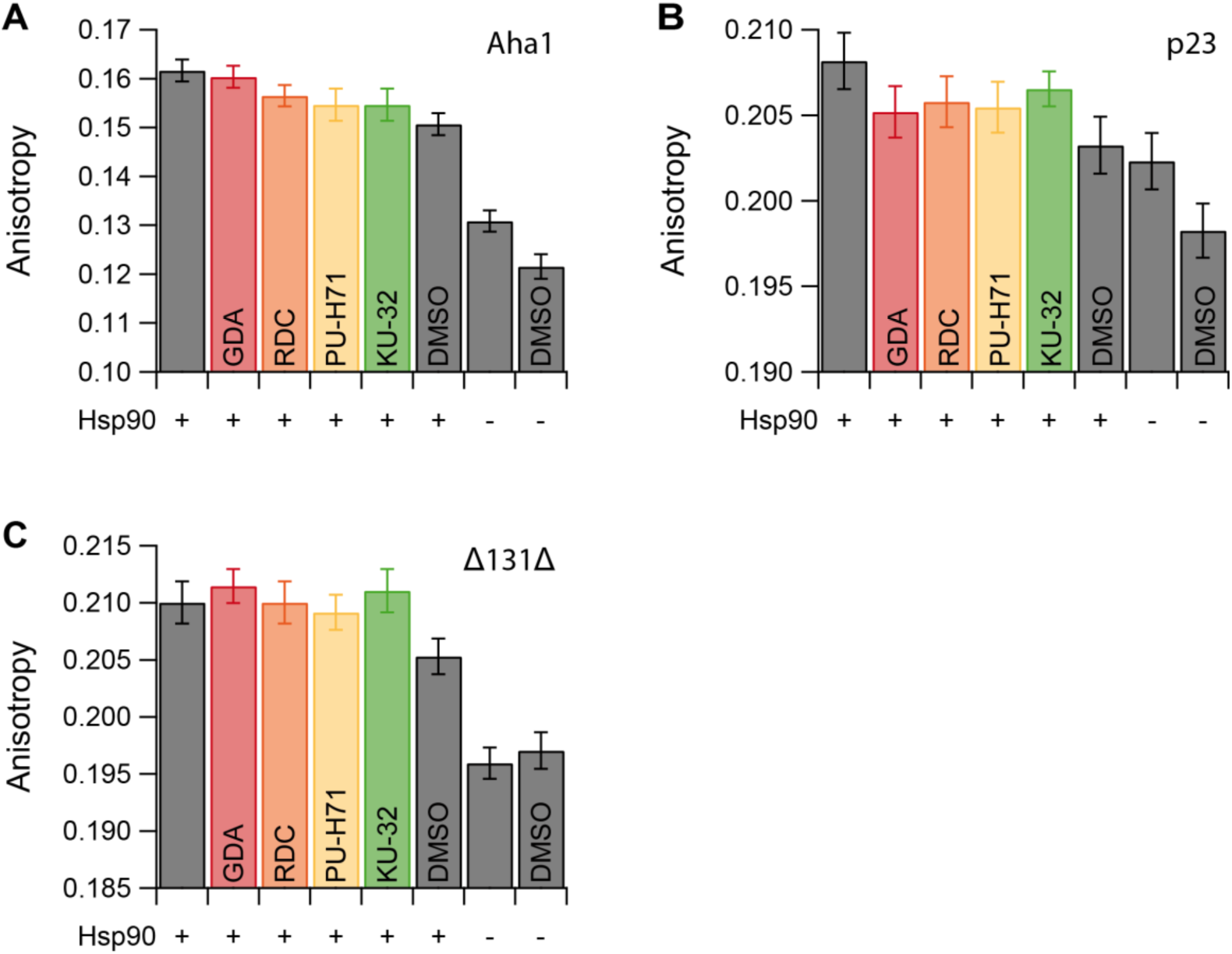
Hsp90 inhibitors show no significant effect on the binding of the co-chaperones Aha1 and p23 or the model client Δ131Δ in fluorescence anisotropy experiments. The fluorescence anisotropy of labeled (**A**) Aha1, (**B**) p23 or (**C**) Δ131Δ is shown in absence or presence of the indicated inhibitor, or DMSO as a control.

## Discussion

To date, the success of Hsp90 inhibitors in clinical trials has been moderate. One reason for this is the lack of a molecular understanding of their precise inhibitory action, preventing a rational drug design. Therefore, we undertook an *in vitro* study covering all four major classes of Hsp90 inhibitors, namely the derivatives of geldanamycin, radicicol, purine and novobiocin. We first determined the complete *in vitro* kinetics of Hsp90’s global opening/closing transitions, under these inhibitor conditions. Interestingly, our results show that the characteristic transitions between globally open and closed conformations are hardly affected, although they are commonly believed to be rate-limiting for Hsp90’s activity (***Röhl et al., 2013***). The observed alterations are neither systematic, nor strong, compared to the effect of AMP-PNP. Thus, it is very unlikely that a direct interference with these conformational changes represents the dominating inhibition mechanism. Similarly, weak effects are found for the interaction with typical cochaperones and a well-characterized model substrate. Therefore, an exclusive interference with one of these interactions is not a likely inhibition mechanism, either.

In addition, our findings help clarify some open points concerning Hsp90’s working principle and inhibition mechanism. First, Hsp90’s ATPase function is still controversially discussed (***Prodromou, 2012, 2016; Schopf et al., 2017; Zierer et al., 2016***). A direct coupling of Hsp90’s characteristic conformational changes to ATP hydrolysis is repeatedly proposed, despite contradicting evidence. If such a direct coupling was the case, competitive ATPase inhibition should ultimately abolish Hsp90’s characteristic conformational changes. On the contrary, the presented results clearly show that such a causality must be dismissed, which supports earlier evidence (***Mickler et al., 2009***) indicating that Hsp90’s characteristic conformational dynamics do not rely on ATPase function. However, it is important to note here that the opposite conclusion does not necessarily hold. I.e. it is still possible that those dynamics are themselves rate-limiting for the slow ATPase rate, although being independent of ATP hydrolysis themselves. Second, Novobiocin derived inhibitors bind to the C-terminal domain, which represents the global hinge of the Hsp90 dimer. Therefore, effects on the large conformational dynamics and associated allosteric regulation were initially expected, but not observed in any of our experiments. Further discussed mechanisms of Novobiocin derivatives include interference with client (***Marcu et al., 2000b***) and/or co-chaperone (namely p23) interaction (***Matts et al., 2011; Yun et al., 2004***). None of these proposed mechanisms withstood our *in vitro* testing.

We like to stress that, although *in vitro* experiments clearly fail to mimic the complex cellular conditions, they remain a valid and crucial hypothesis test. If any of the investigated interactions or conformational changes were directly affected by the inhibitors, it would be detectable *in vitro*, too. In turn, this would have been an ideal *in vitro* test to assess the potency of drug candidates.

Altogether our findings point towards a combination of small interferences, which jointly lead to the observed inhibitory effects. Such combined effects, e.g. on clients and co-chaperones, are difficult to capture in any experiments, because they occur simultaneously and hence they are undetectable in ensemble experiments. In addition, Hsp90’s low-affinity interactions make single molecule experiments, similar to the ones described here, with multiple interaction partners very difficult. As a result, there is no foreseeable reliable *in vitro* test to assess the potency of Hsp90 drug candidates, as neither ATPase activity, nor conformational changes, nor binding of interaction partners are capable to predict the *in vivo* effect.

Nevertheless, our results are the basis for future single molecule experiments in yeast lysate or even live human cell lines to possibly test the potency of drug candidates. We anticipate that our results also help in moving from sheer trial and error to biomedical comprehension of Hsp90’s diverse functions.

## Materials and Methods

### Protein construct preparation

Yeast Hsp90 dimers (UniProtKB: P02829) supplied with a C-terminal coiled-coil motif (kinesin neck region of *Drosophila melanogaster*) were used to avoid dissociation at low concentrations (***Mickler et al., 2009***). Cysteine point mutations allowed specific labeling with donor (D61C) or acceptor (Q385C) fluorophores (see below). Both constructs were cloned into a pET28b vector (Novagen, Merck Biosciences, Billerica, MA). They include an N-terminal His-tag followed by a SUMO-domain for later tag cleavage. The QuikChange Lightning kit (Agilent, Santa Clara, CA) was used to insert an Avitag for specific in vivo biotinylation at the C-terminus of the acceptor construct. *Escherichia coli* BL21star cells (Invitrogen, Carlsbad, CA) were co-transformed with pET28b and pBirAcm (Avidity Nanomedicines, La Jolla, CA) by electroporation (Peqlab, Erlangen, Germany) and expressed according to Avidity’s in vivo biotinylation protocol. The donor construct was expressed in *E. coli* BL21(DE3)cod+ (Stratagene, San Diego, CA) for 3 h at 37°C after induction with 1 mM isopropyl β-D-1-thiogalactopyranoside (IPTG) at OD_600_ = 0.7 in LB_Kana_. A cell disruptor (Constant Systems, Daventry, United Kingdom) was used for lysis. Proteins were purified as published (***Jahn et al., 2014***) (Ni-NTA, tag cleavage, Ni-NTA, anion exchange, size exclusion chromatography). 95% purity was confirmed by SDS-PAGE. Fluorescent labels (Atto550- and Atto647N-maleimide) were purchased from Atto-tec (Siegen, Germany) and coupled to cysteines according to the supplied protocol. Yeast Aha1 (UniProtKB: Q12449) was fluorescently labeled by replacement of S85 with the unnatural amino acid cyclooctyne-lysine (SCO-L-lysine, Sirius Fine Chemicals SiChem GmbH, Bremen, Germany) and coupling with azide-Atto647N. Aha1 was a kind gift of Philipp Wortmann. Yeast p23 (Sba1, UniProtKB: P28707) was expressed as S2C mutant and fluorescently labeled with maleimide-Atto647N. p23 was a kind gift of Johann Thurn. Δ131Δ was expressed as K16C mutant (***Street et al., 2011***) and fluorescently labeled with maleimide-Atto647N. If not stated differently, all chemicals were purchased from Sigma Aldrich.

### Single Molecule FRET measurements

smFRET was measured as previously detailed using a home built TIRF setup (***Schmid et al., 2016***). Hetero-dimers (acceptor + donor) were obtained by 20 min incubation of 1 μM donor and 0.1 μM biotinylated acceptor homodimers in measurement buffer (40 mM HEPES, 150 mM KCl, and 10 mM MgCl_2_, pH7.5) at 47°C. In this way, predominantly biotinylated heterodimers bind to the polyethylene glycol (PEG, Rapp Polymere, Tuebingen, Germany) passivated and neutravidin (Thermo Fisher Scientific, Waltham, MA) coated fluid chamber. Residual homodimers are recognized using alternating laser excitation (ALEX) of donor and acceptor dyes (***Lee et al., 2005***) and excluded from analysis.

All inhibitors were applied at concentrations 100-fold higher than the reported dissociation constant - or the half maximal inhibitory concentration (IC_50_) if the former was not available. The specific concentrations were 100 μM GDA (***Roe et al., 1999***), 1 μM RDC (***Roe et al., 1999***), 12 μM PU-H71 (***He et al., 2006***), 10 μM KU-32 (***Lu et al., 2009***). ADP and AMP-PNP were used at 2 mM.

### Fluorescence anisotropy measurements

Fluorescence anisotropy measurements were performed in measurement buffer. 10 μM Hsp90 with C-terminal coiled-coil (see above) and ± 50 μM inhibitor were added to 200 nM labeled Aha1, 400 nM labeled p23 or 400 nM labeled Δ131Δ. Measurements were performed on a Horiba Fluoro-Max 4 fluorescence spectrometer at 25°C, with excitation at 648 nm (3 nm bandwidth), emission at 660 nm (3 nm bandwidth), 2 s integration time.

## Acknowledgements

We thank G. Chiosis and L. M. Neckers for helpful discussions and providing PU-H71. We thank B. S. J. Blagg for providing KU-32. We thank Philipp Wortmann for Aha1 and Johann Thurn for p23.

This work was funded by the European Research Council through ERC grant agreement no. 681891 and by the German Science Foundation (SFB863, A4).

## References

Ali MMU, Roe SM, Vaughan CK, Meyer P, Panaretou B, Piper PW, Prodromou C, Pearl LH. 2006. Crystal structure of an Hsp90-nucleotide-p23/Sba1 closed chaperone complex. Nature 440:1013–1017. doi: 10.1038/nature04716.

Anyika M, McMullen M, Forsberg LK, Dobrowsky RT, Blagg BSJ. 2016. Development of Noviomimetics as C-Terminal Hsp90 Inhibitors. ACS Medicinal Chemistry Letters 7:67–71. doi: 10.1021/acsmedchemlett.5b00331.

Biamonte MA, van de Water R, Arndt JW, Scannevin RH, Perret D, Lee W-C. 2010. Heat Shock Protein 90: Inhibitors in Clinical Trials. Journal of medicinal chemistry 53:3–17. doi: 10.1021/jm9004708.

Blagg BSJ, Kerr TD. 2006. Hsp90 inhibitors: Small molecules that transform the Hsp90 protein folding machinery into acatalyst for protein degradation. Medicinal Research Reviews 26:310–338. doi: 10.1002/med.20052.

Chiosis G, Timaul MN, Lucas B, Munster PN, Zheng FF, Sepp-Lorenzino L, Rosen N. 2001. A small molecule designed to bind to the adenine nucleotide pocket of Hsp90 causes Her2 degradation and the growth arrest and differentiation of breast cancer cells. Chemistry & Biology 8:289–299. doi: 10.1016/S1074-5521(01)00015-1.

DeBoer C, Meulman PA, Wnuk RJ, Peterson DH. 1970. Geldanamycin, a new antibiotic. The Journal of antibiotics 23:442–447.

Delmotte P, Delmotte-Plaquee J. 1953. A New Antifungal Substance of Fungal Origin. Nature 171:344. doi: 10.1038/171344a0.

Dutta R, Inouye M. 2000. GHKL, an emergent ATPase/kinase superfamily. Trends in Biochemical Sciences 25:24–28. doi: 10.1016/S0968-0004(99)01503-0.

Garg G, Khandelwal A, Blagg BSJ. 2016. Chapter Three - Anticancer Inhibitors of Hsp90 Function: Beyond the Usual Suspects. In: Isaacs J, Whitesell L, eds. Hsp90 in Cancer: Beyond the Usual Suspects. Academic Press.

Ha T. 2001. Single-molecule fluorescence resonance energy transfer. Methods 25:78–86. doi: 10.1006/meth.2001.1217.

He H, Zatorska D, Kim J, Aguirre J, Llauger L, She Y, Wu N, Immormino RM, Gewirth DT, Chiosis G. 2006. Identification of potent water soluble purine-scaffold inhibitors of the heat shock protein 90. Journal of medicinal chemistry 49:381–390.

Hellenkamp B, Wortmann P, Kandzia F, Zacharias M, Hugel T. 2017. Multidomain structure and correlated dynamics determined by self-consistent FRET networks. Nat Meth 14:174–180.

Immormino RM, Kang Y, Chiosis G, Gewirth DT. 2006. Structural and Quantum Chemical Studies of 8-Aryl-sulfanyl Adenine Class Hsp90 Inhibitors. Journal of medicinal chemistry 49:4953–4960. doi: 10.1021/jm060297x.

Jahn M, Rehn A, Pelz B, Hellenkamp B, Richter K, Rief M, Buchner J, Hugel T. 2014. The charged linker of the molecular chaperone Hsp90 modulates domain contacts and biological function. Proc. Natl. Acad. Sci. USA 111:17881–17886. doi: 10.1073/pnas.1414073111.

Khandelwal A, Crowley VM, Blagg BSJ. 2016. Natural Product Inspired N-Terminal Hsp90 Inhibitors: From Bench to Bedside? Medicinal Research Reviews 36:92–118. doi: 10.1002/med.21351.

Lee NK, Kapanidis AN, Wang Y, Michalet X, Mukhopadhyay J, Ebright RH, Weiss S. 2005. Accurate FRET Measurements within Single Diffusing Biomolecules Using Alternating-Laser Excitation. Biophysical Journal 88:2939–2953.

Li J, Richter K, Reinstein J, Buchner J. 2013. Integration of the accelerator Aha1 in the Hsp90 co-chaperone cycle 20:326 EP -. doi: 10.1038/nsmb.2502.

Lu Y, Ansar S, Michaelis ML, Blagg BSJ. 2009. Neuroprotective activity and evaluation of Hsp90 inhibitors in an immortalized neuronal cell line. Bioorganic & Medicinal Chemistry 17:1709–1715. doi: 10.1016/j.bmc.2008.12.047.

Marcu MG, Chadli A, Bouhouche I, Catelli M, Neckers LM. 2000a. The Heat Shock Protein 90 Antagonist Novobiocin Interacts with a Previously Unrecognized ATP-binding Domain in the Carboxyl Terminus of the Chaperone. Journal of Biological Chemistry 275:37181–37186. doi: 10.1074/jbc.M003701200.

Marcu MG, Schulte TW, Neckers L. 2000b. Novobiocin and Related Coumarins and Depletion of Heat Shock Protein 90-Dependent Signaling Proteins. Journal of the National Cancer Institute 92:242–248. doi: 10.1093/jnci/92.3.242.

Matts RL, Dixit A, Peterson LB, Sun L, Voruganti S, Kalyanaraman P, Hartson SD, Verkhivker GM, Blagg BSJ. 2011. Elucidation of the Hsp90 C-Terminal Inhibitor Binding Site. ACS Chemical Biology 0. doi: 10.1021/cb200052x.

McLaughlin SH, Sobott F, Yao Z-p, Zhang W, Nielsen PR, Grossmann JG, Laue ED, Robinson CV, Jackson SE. 2006. The Co-chaperone p23 Arrests the Hsp90 ATPase Cycle to Trap Client Proteins. Journal of Molecular Biology 356:746–758. doi: 10.1016/j.jmb.2005.11.085.

Mickler M, Hessling M, Ratzke C, Buchner J, Hugel T. 2009. The large conformational changes of Hsp90 are only weakly coupled to ATP hydrolysis. Nat Struct Mol Biol 16:281–286. doi: 10.1038/nsmb.1557.

Neckers L, Workman P. 2012. Hsp90 Molecular Chaperone Inhibitors: Are We There Yet? Clinical Cancer Research 18:64–76. doi: 10.1158/1078-0432.CCR-11-1000.

Prodromou C. 2012. The ‘active life’ of Hsp90 complexes. Biochimica et Biophysica Acta (BBA) - Molecular Cell Research 1823:614–623. doi: 10.1016/j.bbamcr.2011.07.020.

Prodromou C. 2016. Mechanisms of Hsp90 regulation. Biochemical Journal 473:2439–2452. doi: 10.1042/BCJ20160005.

Rodina A, Vilenchik M, Moulick K, Aguirre J, Kim J, Chiang A, Litz J, Clement CC, Kang Y, She Y, Wu N, Felts S, Wipf P, Massague J, Jiang X, Brodsky JL, Krystal GW, Chiosis G. 2007. Selective compounds define Hsp90 as a major inhibitor of apoptosis in small-cell lung cancer. Nat Chem Biol 3:498–507. doi: 10.1038/nchembio.2007.10.

Roe SM, Prodromou C, O’Brien R, Ladbury JE, Piper PW, Pearl LH. 1999. Structural Basis for Inhibition of the Hsp90 Molecular Chaperone by the Antitumor Antibiotics Radicicol and Geldanamycin. Journal of medicinal chemistry 42:260–266. doi: 10.1021/jm980403y.

Röhl A, Rohrberg J, Buchner J. 2013. The chaperone Hsp90: Changing partners for demanding clients. Trends in Biochemical Sciences 38:253–262. doi: 10.1016/j.tibs.2013.02.003.

Sattin S, Tao J, Vettoretti G, Moroni E, Pennati M, Lopergolo A, Morelli L, Bugatti A, Zuehlke A, Moses M, Prince T, Kijima T, Beebe K, Rusnati M, Neckers L, Zaffaroni N, Agard DA, Bernardi A, Colombo G. 2015. Activation of Hsp90 Enzymatic Activity and Conformational Dynamics through Rationally Designed Allosteric Ligands. Chemistry - A European Journal 21:13598–l13608. doi: 10.1002/chem.201502211.

Schmid S, Götz M, Hugel T. 2016. Single-Molecule Analysis beyond Dwell Times: Demonstration and Assessment in and out of Equilibrium. Biophysical Journal 111:1375–1384. doi: 10.1016/j.bpj.2016.08.023.

Schopf FH, Biebl MM, Buchner J. 2017. The HSP90 chaperone machinery. Nat Rev Mol Cell Biol 18:345–360.

Sidera K, Patsavoudi E. 2014. HSP90 Inhibitors: Current Development and Potential in Cancer Therapy. Recent Patents on Anti-Cancer Drug Discovery 9:1–20. doi: 10.2174/15748928113089990031.

Southworth DR, Agard DA. 2008. Species-Dependent Ensembles of Conserved Conformational States Define the Hsp90 Chaperone ATPase Cycle. Molecular Cell 32:631–640.

Street TO, Lavery LA, Agard DA. 2011. Substrate Binding Drives Large-Scale Conformational Changes in the Hsp90 Molecular Chaperone. Molecular Cell 42:96–105. doi: 10.1016/j.molcel.2011.01.029.

Taipale M, Jarosz DF, Lindquist S. 2010. HSP90 at the hub of protein homeostasis: Emerging mechanistic insights. Nat Rev Mol Cell Biol 11:515–528. doi: 10.1038/nrm2918.

Taldone T, Patel HJ, Bolaender A, Patel MR, Chiosis G. 2014. Protein chaperones: A composition of matter review (2008 - 2013). Expert Opinion on Therapeutic Patents 24:501–518. doi: 10.1517/13543776.2014.887681.

Yun B-G, Huang W, Leach N, Hartson SD, Matts RL. 2004. Novobiocin Induces a Distinct Conformation of Hsp90 and Alters Hsp90-Cochaperone-Client Interactions. Biochemistry 43:8217–8229. doi: 10.1021/bi0497998.

Zierer BK, Rubbelke M, Tippel F, Madl T, Schopf FH, Rutz DA, Richter K, Sattler M, Buchner J. 2016. Importance of cycle timing for the function of the molecular chaperone Hsp90. Nat Struct Mol Biol 23:1020–1028.

Zuehlke AD, Moses MA, Neckers L. 2018. Heat shock protein 90: Its inhibition and function. Philosophical Transactions of the Royal Society B: Biological Sciences 373. doi: 10.1098/rstb.2016.0527.

